# Acute stress blunts prediction error signals in the dorsal striatum during reinforcement learning

**DOI:** 10.1101/2021.02.11.430640

**Authors:** Joana Carvalheiro, Vasco A. Conceição, Ana Mesquita, Ana Seara-Cardoso

## Abstract

Reinforcement learning, which implicates learning from the rewarding and punishing outcomes of our choices, is critical for adjusted behaviour. Acute stress seems to affect this ability but the neural mechanisms by which it disrupts this type of learning are still poorly understood. Here, we investigate whether and how acute stress blunts neural signalling of prediction errors during reinforcement learning using model-based functional magnetic resonance imaging. Male participants completed a well-established reinforcement learning task involving monetary gains and losses whilst under stress and control conditions. Acute stress impaired participants’ behavioural performance towards obtaining monetary gains, but not towards avoiding losses. Importantly, acute stress blunted signalling of prediction errors during gain and loss trials in the dorsal striatum — with subsidiary analyses suggesting that acute stress preferentially blunted signalling of positive prediction errors. Our results thus reveal a neurocomputational mechanism by which acute stress may impair reward learning.

## Introduction

The ability to gradually learn from the consequences of our actions, to make choices that lead to the best possible outcomes, is crucial for adaptive behaviour. This ability seems to be affected by situational factors, including those that are ubiquitous in modern everyday life. A growing body of evidence suggests that reward learning is impaired by acute stress^1–9^, although the evidence for an impairing effect of acute stress on punishment learning is less robust^7,9,10^. Yet, surprisingly little is known about the neural mechanisms that underlie the impairing effects of acute stress on reinforcement learning. Given the pervasiveness of stress, characterising the neural mechanisms by which acute stress affects our ability to learn from obtained rewards and from avoided punishments is relevant to understand the effects of stress in everyday life, and it might offer important insight into the development of treatments for individuals with stress-related clinical disorders. Here, we use behavioural and model-based functional magnetic resonance imaging (fMRI)^11^ data to investigate the impact of acute stress on reinforcement learning and the underlying neurocomputational mechanisms.

Reinforcement-learning theory provides a powerful neurocomputational framework to understand how individuals learn to maximise rewards and minimise punishments^12,13^. According to reinforcement-learning theory, individuals gradually learn to select more and more often the actions that optimise reinforcements in a given context by learning the values of the executed actions^12,13^. Prediction errors — the difference between an experienced and an expected outcome — are used to progressively update the values of the executed actions driving gradual learning^12–14^. Positive prediction errors indicate that outcomes are better than expected, and negative prediction errors indicate that outcomes are worse than expected^14^. Prediction errors can therefore be used to learn which actions are advantageous or disadvantageous. For example, when an action results in an outcome that is better than expected, a positive prediction error occurs, and the value of the action is increased, leading to increased likelihood of selecting that action in the future. Prediction error signals are thought to be encoded in the phasic activity of dopamine neurons^14^. Extant evidence indicates that brain areas with dense dopaminergic projections, such as the dorsal striatum and the nucleus accumbens, show activity correlated with prediction errors^15–17^ and that prediction error signals in the dorsal striatum correlate with behavioural performance efficacy in a reward-based task^18^. Indeed, the striatal dopaminergic system seems to be critical for prediction-error-based reward learning^13,15,19,20^.

The striatal dopaminergic system also seems to be particularly sensitive to acute stress^21–23^. Acute stress elicits a myriad of physiological and functional changes in the brain in response to perceived adverse changes in the environment^24–26^, including increased dopamine release in the striatum^21–23,25–28^. Specifically, studies with non-human male animals suggest that acute stress increases aberrant spontaneous phasic-dopamine release^29–31^. Such exaggerated, aberrant spontaneous dopamine release is thought to blunt adaptive phasic dopamine responses that signal positive prediction errors^32–34^ and, more tentatively, negative prediction errors^32^. Thus, stress-induced dopamine aberrant release may lead to impairments in reward learning, and more speculatively in punishment learning. Indeed, extant behavioural data suggest that acute stress impairs reward learning^1–9,35^, whereas the impact of stress on punishment learning remains more equivocal^7,9,10,35^.

Extant neural evidence on how acute stress directly affects prediction error signals in the human striatum during reward learning is still scarce^8,36,37^, but indirect neural evidence indicates that women who show the greatest increase in interleukin-6 (an inflammatory marker) in response to a stressor also show the greatest reduction in signalling of prediction errors in the nucleus accumbens during reinforcement learning^37^. Moreover, we previously showed, using computational modelling, that acute stress decreases the learning rate for positive prediction errors (i.e., how quickly better-than-expected outcomes are integrated over time)^7^, which seems to be consistent with the idea that acute stress might impair reward learning by blunting neural signalling of prediction errors.

In this study, we aimed to investigate the impact of acute stress on striatal prediction error signalling during reinforcement learning. As mentioned above, extant literature suggests that acute stress disrupts reward learning to a larger extent than punishment learning. Thus, given the putative impact of acute stress on aberrant phasic-dopamine release, and the role of adaptive phasic-dopamine responses on prediction error signalling, we predicted that 1) acute stress would impair reward learning and, relatedly, that 2) acute stress would blunt prediction error signals in the striatum during reward learning. Additionally, given that striatal dopamine prediction error signals are also implicated in punishment learning^38–40^, we explored whether and how acute stress would impact punishment learning. Finally, we explored whether acute stress would preferentially blunt positive or negative prediction error signals during reward and punishment learning.

Thirty-seven male participants completed an adapted version of a well-established reinforcement-learning task involving monetary gains and losses^15^ inside the MRI scanner, whilst under acute stress and control conditions (Fig. 1). This reinforcement-learning task disentangles reward from punishment learning and has been used to assess fluctuations in dopamine-prediction errors signals; using this task, combined with pharmacological manipulations of the dopaminergic system, Pessiglione et al.^15^ showed that dopamine-related drugs modulate prediction errors expressed in the striatum during reward (but not during punishment) learning. In each trial of the reinforcement-learning task, participants were asked to choose between two abstract visual stimuli to maximise payoffs. Each pair of stimuli was associated with a valence: one pair of stimuli was associated with gains (0.5€ or no change), a second pair was associated with losses (−0.5€ or no change), and a third pair was associated with neutral, or non-financial outcomes (look at a 0.5€ coin or no change). The outcome probabilities were reciprocally 0.75 and 0.25 for the two stimuli in each of the three pairs (Fig. 1).

**Fig. 1.**
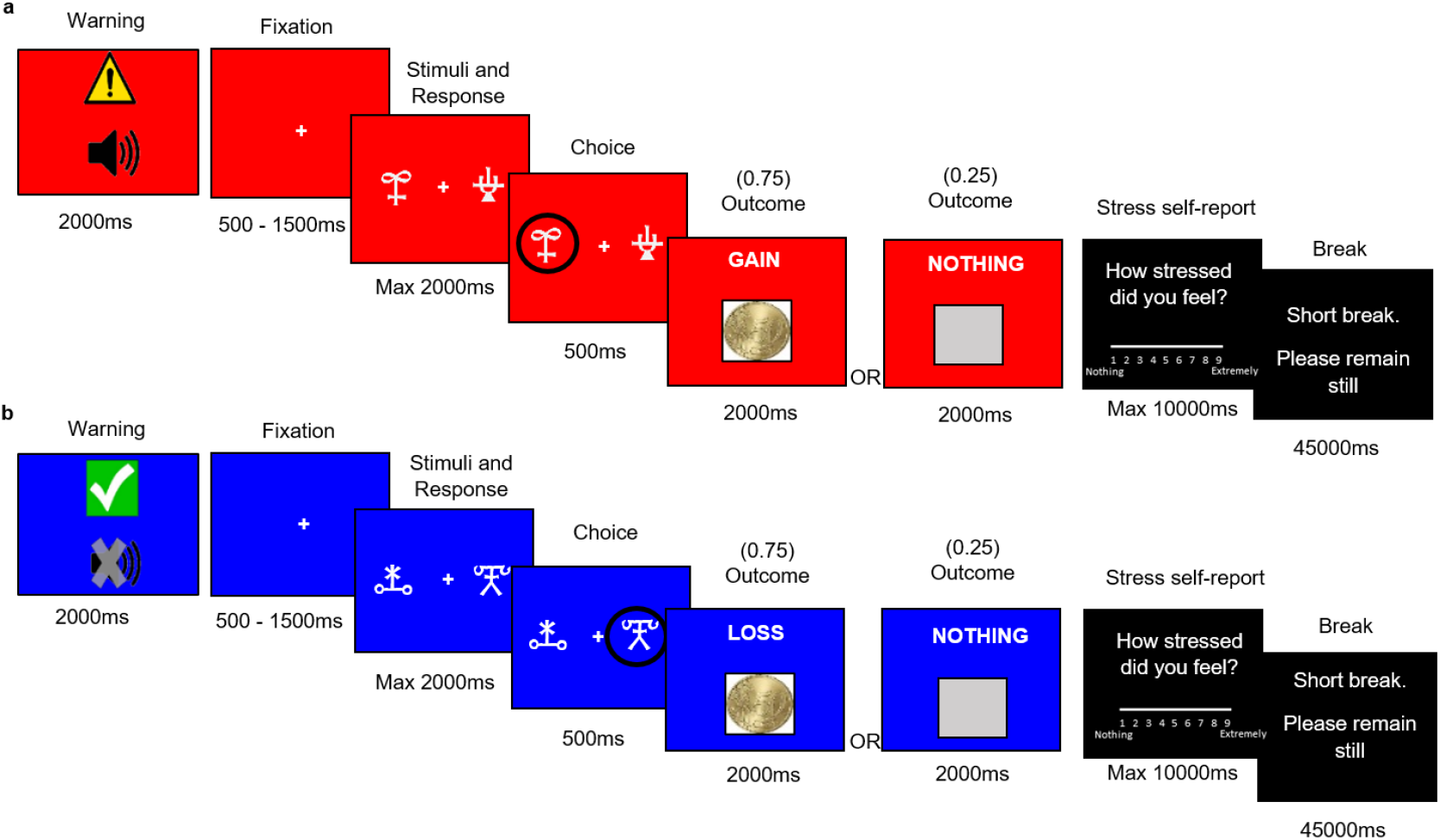
Reinforcement-learning task. Inside the scanner, participants chose between two abstract visual stimuli and observed the outcome of their choice, whilst under acute stress and under control conditions. **(a)** In the depicted stress-condition trial, the chosen stimulus was associated with a probability of 0.75 of winning 0.5€ and with a probability of 0.25 of winning nothing. The other (not chosen) stimulus was associated with a reciprocate probability of 0.75 of winning nothing and a 0.25 probability of winning 0.5€. **(b)** In the depicted control-condition example, the chosen stimulus was associated with a probability of 0.75 of losing 0.5€ and with a probability of 0.25 of losing nothing. The other (not chosen) stimulus was associated with a reciprocate probability of 0.75 of losing nothing and with a 0.25 probability of losing 0.5€. Participants completed a total of four blocks (two stress and two control blocks), consisting of an alternation between stress and control blocks. To assess stress responses, self-reported stress levels were collected at the end of each block.

Participants completed the task in four blocks. In half of the blocks, participants were exposed to an uncontrollable stressor, a constant alarm (i.e., stress condition), which was previously shown to be effective in increasing self-reported stress levels and skin conductance responses rate^7^. These blocks were alternated with blocks without the stressor (i.e., control condition). To check the success of the acute-stress manipulation, we collected self-reported stress levels at the end of each block. Participants who reported to be non-responsive to the stress manipulation (i.e., who did not report higher stress levels in the stress condition than in the control condition) were excluded from the main analyses. This resulted in a final pool of 23 participants for behavioural and neuroimaging data analyses. For completeness, we also analysed the data from the total sample, which yielded findings for the impact of acute stress on reward learning consistent with those from the analyses of the aforementioned subsample of interest (see the Supplementary Information for analyses and results concerning the total sample).

To assess whether acute stress impaired reward learning, we inspected the impact of acute stress on task performance. Next, to examine whether and how acute stress blunted signalling of prediction errors in the striatum, we used trial-wise prediction errors, estimated by a well-established reinforcement-learning model^41^, as parametric modulators of striatal — dorsal striatum and nucleus accumbens — activity at the time of the outcomes in gain (i.e., reward learning) and loss (i.e., punishment learning) trials^15^.

## Results

### Behavioural analyses

#### Manipulation check

First, we computed the difference in self-reported stress levels between the stress and control conditions in the total sample (n = 37). Twenty-three participants reported higher stress levels in the stress condition than in the control condition (Fig. 2a). Then, we conducted analyses of variance (ANOVAs) with condition (stress and control) and block (1 and 2) as within-subject factors in those 23 participants (see Supplementary Information for manipulation-check analyses of the total sample). Self-reported stress levels differed significantly between conditions (*F*_1,22_= 69.28, *p* < 0.001, *ƞ*^*2*^ = 0.76) (Fig. 2b), but there was no main effect of block (*F*_1,22_ = 0.008, *p* = 0.93, *ƞ*^*2*^ = 0) and the condition × block interaction was also non-significant (*F*_1,22_ = 1.21, *p* = 0.28, *ƞ*^*2*^ = 0.052). This suggests that self-reported stress levels increased with the acute stress manipulation and remained stable across blocks within each condition for these participants. Participants whose self-reported levels of stress did not increase with the acute-stress induction were excluded from the following analyses (but see Supplementary Information for total sample analyses).

**Fig. 2.**
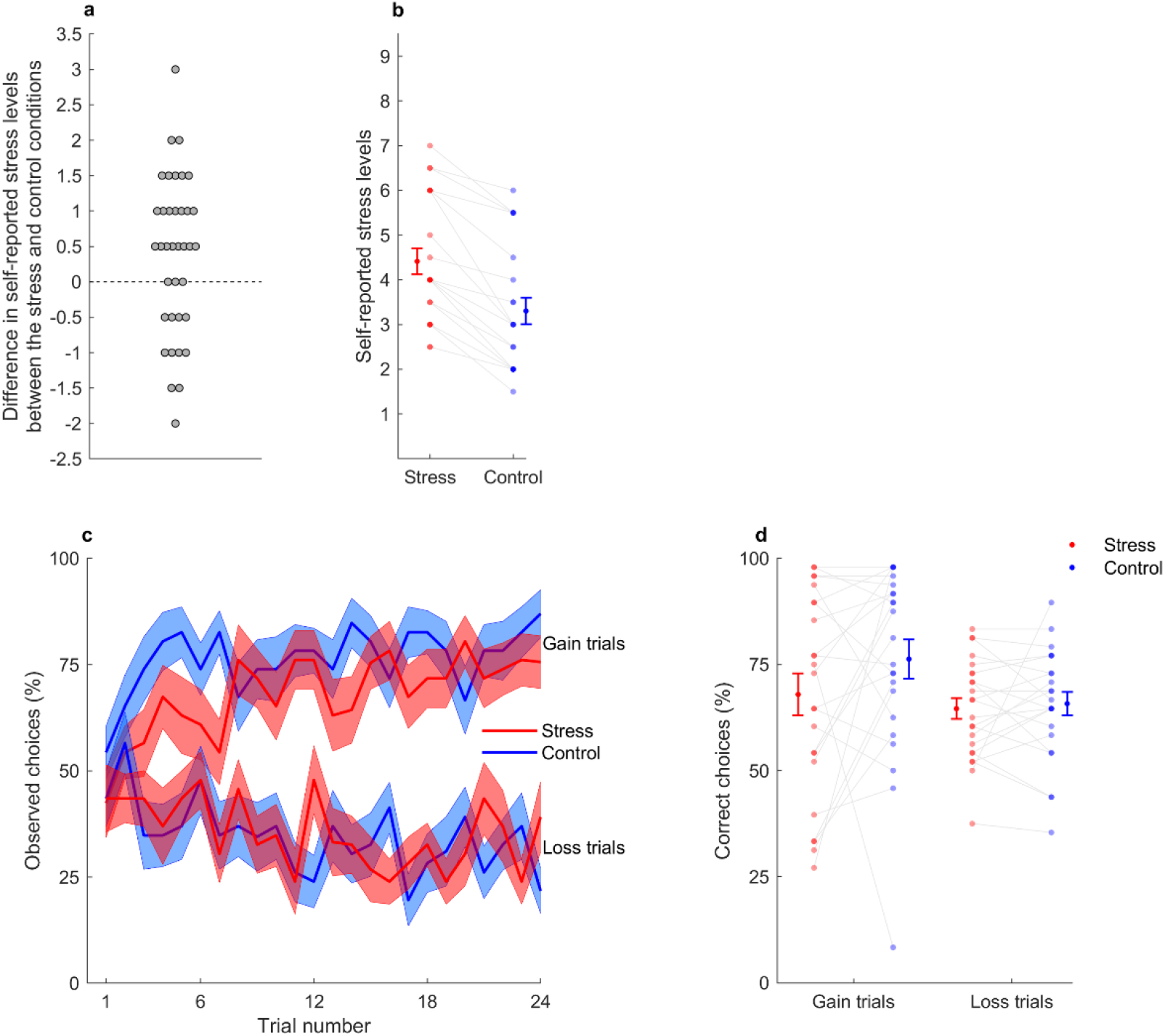
Manipulation check and task performance. **(a)** Difference in self-reported stress levels between the stress and control conditions (averaged across blocks). Each grey dot represents a participant (n = 37). Participants who reported higher stress levels in the stress than in the control condition correspond to the dots above the horizontal dashed line (n = 23). **(b)** Self-reported stress levels in stress (red) and control (blue) conditions (averaged across blocks) in the pool of participants that reported higher stress levels in the stress than in the control condition (n = 23). **(c)** Learning curves represent the trial-by-trial percentage of participants (n = 23) who chose the correct gain stimulus (associated with a probability of 0.75 of winning 0.5€; upper part of the graph) and the incorrect loss stimulus (associated with a probability of 0.75 of losing 0.5€; lower part of the graph), in the stress and control conditions. Each central line represents the mean and each filled area the ± standard error of the mean. **(d)** Percentage of correct choices per participant (n = 23) in gain and loss trials, across the stress and control conditions (averaged across blocks). In graphs b and d, connected dots represent data points from the same participant, and more transparent (opaque) dots represent less (more) overlapping data points; error bars displayed on the sides of those scatter plots indicate the mean ± standard error of the mean.

#### Task performance

To assess whether acute stress would blunt reward learning, we examined the impact of acute stress on choice performance during the reinforcement-learning task (Fig. 2c). We used a generalized linear mixed-effects (glme) model, which accounted for the binomial distribution of the trial-by-trial data (correct or incorrect responses). The glme model included condition (stress or control), valence (gain or loss), block number (1 or 2), trial number (1 to 24), and the interaction of interest (condition × valence) as predictor variables. We found a significant condition × valence interaction (*β* = -0.39, *p* = 0.0038, 95% CI = [-0.66, -0.13]) (Fig. 2d). Planned post-hoc analyses showed that under stress, comparatively to the control condition, participants performed significantly worse when learning to obtain gains (*F*_1, 4400_ = 20.23, *p* < 0.001), but not when learning to avoid losses (*F*_1, 4400_ = 0.32, *p* = 0.57). In sum, acute stress selectively impaired choice performance towards monetary gains during the reinforcement-learning task.

### fMRI analyses

To examine the impact of acute stress on prediction error signalling in the striatum during reinforcement learning, we generated a primary fMRI general linear model that included prediction errors as parametric modulators of BOLD response in the striatum (dorsal striatum and nucleus accumbens) at the time of the outcomes in gain and loss trials, in the stress and control conditions (see Methods and Supplementary Fig. 5 for further details on the primary general linear model). Prediction errors were estimated in the stress and control conditions using a reinforcement-learning model that has been extensively used to investigate the behavioural and neural impact of pharmacological manipulations and genetic variations in the dopaminergic system in humans^41–46^ (see Supplementary Information for computational modelling methods and results). Parametric analyses incorporating prediction errors allow a more precise estimation of how brain activity fluctuates during learning compared to examination of outcome-associated activation alone^11^.

As expected, we observed that BOLD response in the dorsal striatum and nucleus accumbens — regions consistently shown to respond to unexpected rewards and punishments^15,47^ — scaled parametrically with the magnitude of prediction errors at the time of the outcomes in gain and loss trials in the stress and/or control conditions. Specifically, we identified a positive parametric modulation of prediction errors in the dorsal striatum bilaterally (i.e., the magnitude of the prediction errors correlated positively with BOLD response in this region on a trial-by-trial basis), during gain and loss trials, both in the stress and control conditions [all *Z* > 4.12, *p* < 0.05, voxel-level small-volume family-wise error corrected (SVC-FWE)]. We also found a positive parametric modulation of prediction errors in the nucleus accumbens during gain trials, both in the stress and control conditions, and during loss trials in the control condition (all Z > 3.63, *p* < 0.05, SVC-FWE; see Supplementary Table 1 for whole-brain and all SVC-FWE results).

After confirming that striatal activity scaled parametrically with the magnitude of prediction errors, we inspected whether acute stress affected prediction error signals in the striatum. To examine whether acute stress would blunt signalling of prediction errors in the striatum during reward learning, first we tested the main effect of stress using the control > stress contrast for the parametric modulators of prediction errors at the time of the outcomes delivered across gain and loss trials in each condition. A significant main effect would mean that acute stress decreased prediction error signals across gain and loss trials. Second, we tested the contrast for the condition (stress or control) × valence (gain or loss) interaction. A significant interaction would mean that acute stress affected prediction error signals differently in gain and loss trials.

The contrast control > stress showed a significant main effect of stress on the parametric modulation of prediction errors in the dorsal striatum ([*x* = 32, *y* = 0, *z* = 12], *Z* = 4.08, *k* =26, *p* = 0.040, SVC-FWE) (Fig.3a), meaning that prediction error signals were decreased in the stress condition compared with the control condition, both in gain and loss trials (Fig. 3b). Confirmatory one sample *t*-tests comparing the parameter estimates (i.e., the regression slopes from the primary general linear model) extracted from the identified dorsal striatum cluster against zero, indicated that the parametric modulation of prediction errors was significantly higher than zero in the control condition, both for gain and loss trials (both *p* < 0.023), but not in the stress condition (both *p* > 0.33) (Fig. 3b). We did not observe any significant responses for the parametric modulation of prediction errors in the nucleus accumbens for the control > stress contrast (nor for the inverse contrast stress > control).

**Fig. 3.**
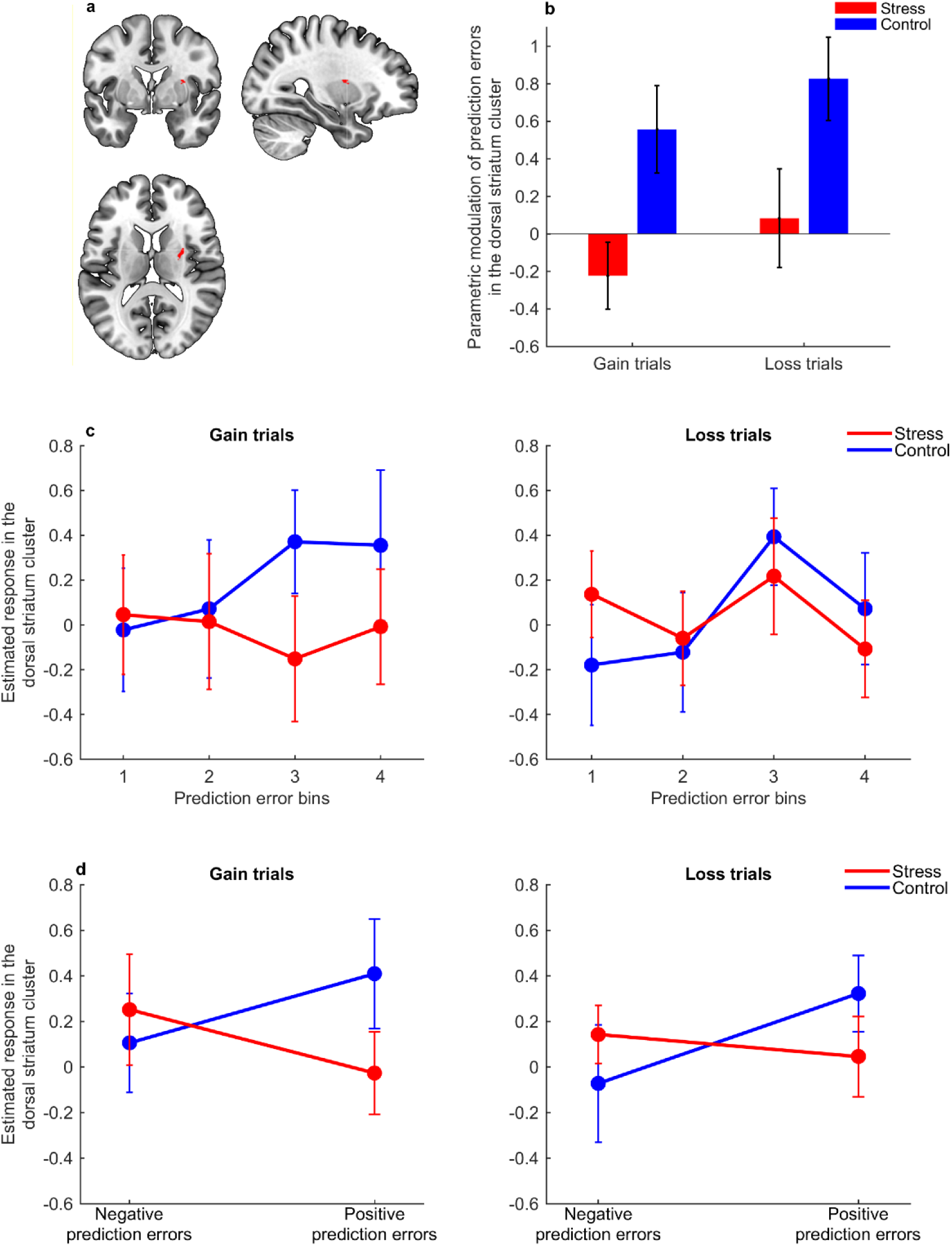
Effects of acute stress on prediction error signalling in the dorsal striatum. **(a)** Cluster in the dorsal striatum where the modulation of prediction errors at the time of the outcome was significantly decreased in the stress condition compared with the control condition (SVC-FWE, *p* < 0.05). **(b)** Bars depict parameter estimates (i.e., regression slopes) for the BOLD response at the dorsal striatum cluster [from (a)] modulated by trial-by-trial prediction errors, in gain and loss trials, across the stress (red) and control (blue) conditions (n = 23). **(c, d)** Graphs represent the modulation of BOLD response by prediction errors at the time of the outcome in gain (left) and loss (right) trials, in the dorsal striatum cluster identified in the primary general linear model [from (a)], in the stress (red) and control (blue) conditions (n = 23). BOLD response estimates from the dorsal striatum cluster were extracted for each participant. Error bars indicate the mean ± standard error of the mean. In **(c)**, data for illustrative graphs were derived from a subsidiary model where trials were divided into quartiles of magnitude of prediction errors (with the lowest and highest magnitudes corresponding to bins 1 and 4, respectively). In **(d)**, the plotted data were obtained from a second subsidiary model where trials were divided into negative and positive prediction errors.

For the contrast that tested the condition × valence interaction, we did not find any significant activations in the dorsal striatum nor in the nucleus accumbens, indicating that the effect of acute stress on prediction error signals was not dependent on the valence of the trial. This non-significant interaction, together with the significant main effect described above, indicates that acute stress blunted prediction errors in the dorsal striatum both in gain and loss trials.

To illustrate the effect of acute stress on the parametric modulation of prediction errors in the dorsal striatum, we conducted a subsidiary general linear model from which we extracted estimates of BOLD response in the dorsal striatum cluster identified in the primary general linear model (cluster represented in Fig. 3a) across trials of different categories of prediction error magnitudes. Specifically, in this subsidiary model, trials from each condition (i.e., stress and control) and valence (i.e., gain and loss) were further divided into four bins corresponding to quartiles of magnitude of prediction errors (see Supplementary Table 2 for median and boundaries of each bin). Parameter estimates of BOLD response at the outcome onset were extracted from the dorsal striatum cluster for each subject (see Methods for full description) and plotted to illustrate the variation in the BOLD response in the dorsal striatum cluster along the magnitude of prediction errors. The blunting main effect of stress (both in gain and loss trials) on prediction error signals in the dorsal striatum seemed to be mostly driven by decreased signalling of prediction errors of higher magnitude (Fig. 3c). It is important to note that the 1^st^ bin and 4^th^ bins roughly correspond to negative and positive prediction errors, respectively (Supplementary Table 2), which suggests that acute stress mostly decreased positive prediction error signals.

To further explore whether acute stress had preferentially blunted signalling of positive prediction errors during reward and punishment learning, we conducted an additional parametric modulation model similar to the primary general linear model, but this time we modelled positive and negative prediction errors separately for either gain or loss trials. We did not find any parametric modulation of the dorsal striatum or nucleus accumbens response by positive nor negative prediction errors, very possibly due to reduced variance within each parametric modulator. Therefore, and for completeness, we performed a second subsidiary general linear model (see Methods for full description). In this subsidiary model, trials from each condition (i.e., stress or control) and valence (i.e., gain or loss) were further divided according to the prediction error valence (i.e., positive or negative). We extracted estimates of BOLD response in the dorsal striatum cluster (cluster represented in Fig. 3a) at the outcome onset, when positive or negative prediction errors occurred, and performed an ANOVA on those estimates derived from the subsidiary model. We found a significant condition × prediction error valence interaction (*F*_1,22_ = 8.84, *p* = 0.007, *η*^2^ = 0.29) (Fig. 3d), indicating that acute stress differently affected positive and negative prediction errors, both in gain and loss trials (the condition × prediction error valence × trial valence interaction was non-significant, *F*_1,22_ = 0.047, *p* = 0.83, *η*^2^ = 0.0020). Post-hoc planned comparisons were non-significant, but inspection of effect sizes suggested that acute stress decreased positive prediction error signals (paired t-tests in gain trials: *t*_22_ = -1.36, p = 0.19, Cohen’s *d* = -0.28; in loss trials: *t*_22_ = -1.12, p = 0.27, Cohen’s *d* = -0.23) to a larger extent than negative prediction errors (paired t-tests in gain trials: *t*_22_= 0.48, *p* = 0.64, Cohen’s *d* = 0.10; in loss trials: *t*_22_ = 0.86, *p* = 0.40, Cohen’s *d* = 0.18) (Fig. 3d), in line with the previous subsidiary analysis of prediction error bins depicted in Fig. 3c.

In sum, we found that the BOLD response in the dorsal striatum scaled parametrically with the magnitude of prediction errors during reward and punishment learning, both under acute stress and control conditions. More importantly, we found that signalling of prediction errors in the dorsal striatum was blunted by acute stress, with subsidiary data suggesting that acute stress mostly decreased positive prediction errors signals.

## Discussion

Previous studies have found that acute stress impacts reinforcement learning. Yet, the mechanisms that underlie the impact of acute stress on reinforcement learning are still poorly understood. Acute stress is ubiquitous in modern day-to-day life. Understanding whether and how acute stress influences our ability to learn from the rewarding and punishing outcomes of our choices might be important for the development of interventions that target the debilitating effects of acute stress. In this study, we investigated whether and how acute stress impacts reinforcement learning using behavioural and model-based fMRI data. Acute stress alters striatal dopaminergic functioning^21–23,28,30,31^, and striatal dopaminergic functioning plays a key role in signalling of prediction errors^13,15,19,20^ — the result of a positive or negative difference between obtained and expected outcomes — which drive reward and punishment learning. Thus, we set out to test whether acute stress impaired reward learning by blunting prediction error signals in the striatum and further explored the putative impact of acute stress on punishment learning.

In line with extant literature^1–6,8,9^, we replicated our previous finding that acute stress impairs performance towards rewards in a reinforcement-learning task^7^. Importantly, we also found that such behavioural impairment was accompanied by blunted signalling of prediction errors in the dorsal striatum. Additionally, even though we did not find evidence of the impact of acute stress on punishment learning at the behavioural level, neural data indicated that acute stress also blunted prediction error signals in the dorsal striatum during punishment learning. Subsidiary analyses indicated that acute stress may blunt preferentially positive prediction error signals, which might explain the differential impact of acute stress in reward and punishment learning.

Our finding that acute stress blunted signalling of prediction errors — and mostly positive prediction errors — in the dorsal striatum during reward learning is consistent with a neurobiological framework of stress-induced dopamine disruptions. Prediction errors are encoded in phasic activity of dopaminergic neurons^14,19,20^. Phasic bursts of dopaminergic neurons are thought to adaptively encode positive prediction errors, whereas dopamine dips have been associated with the adaptive encoding of negative prediction errors^14,19^. However, phasic-dopamine responses do not seem to be always adaptive, and there is evidence that dopamine can be phasically released in an aberrant spontaneous manner^32,48–50^. Relatedly, studies with non-human male animals suggest that acute stress induces aberrant spontaneous dopamine release^29–31^. It is therefore possible that, if acute stress increases aberrant spontaneous phasic-dopamine release, then phasic dopamine release that signals positive prediction errors following burst firing of dopaminergic neurons is less easily differentiated from background fluctuations in dopamine levels, resulting in a low signal to noise ratio^48,50^. In addition, if there is increased aberrant spontaneous release of dopamine, then less dopamine may be available to be released from dopaminergic neurons when positive prediction errors occur^48^. Thus, stress-induced aberrant dopamine release may indirectly or directly blunt positive prediction errors that signal unexpected rewards^32–34^, resulting in impaired reward learning. Furthermore, we previously showed, using computational modelling, that acute stress decreases the learning rate for positive prediction errors^7^, which is in striking agreeement with our neuroimaging data. Our finding that acute stress blunts prediction errror signals — and preferentially positive prediction errors — in the dorsal striatum during reward learning is thus consistent with a neurobiological framework of stress-induced striatal dopaminergic disturbances and might explain why individuals have difficulties in updating their behaviour in response to unexpected rewards when under acute stress.

In this study, we also explored how acute stress affected punishment learning, although the behavioural evidence for an effect of acute stress on punishment learning seems less robust than for an effect of stress on reward learning^35^. In line with our previous work, using the same stress manipulation and an adapted version of the same reinforcement-learning task in an independent sample^7^, we found no evidence of a behavioural impairment of acute stress on punishment learning. This suggests that punishment learning may not be affected by acute stress to the same extent as reward learning is. However, our neuroimaging data showed a main effect of stress on the signalling of prediction errors in the dorsal striatum, indicating that acute stress also decreased prediction error signals during punishment learning. Further subsidiary analyses suggest that the main effect of stress on prediction errors seem to be mostly explained by decreased signalling of positive prediction errors. It is thus possible that acute stress compromises the ability to use dopamine phasic bursts that signal positive prediction errors but not to use dopamine dips that encode negative prediction errors. Indeed, empirical evidence suggests that aberrant spontaneous dopamine release decreases striatal adaptive phasic dopamine responses that signal positive prediction errors^33,34^, and, only more speculatively, negative prediction errors^32^. Although positive prediction errors can also occur during punishment learning, in simple reinforcement-learning tasks, such as ours, punishment learning seems to be largely driven by negative prediction errors^38^. Consequently, stress-induced disruptions in positive prediction errors might not necessarily be reflected in impaired learning from punishments. Finally, non-dopaminergic mechanisms may also be involved in punishment learning^51,52^, which might partially explain why previous studies using similar reinforcement-learning tasks also did not find significant effects of pharmacological manipulations of the dopaminergic system on punishment learning^15,53^.

We found evidence for stress-induced blunted prediction errors in the dorsal striatum, but not in the nucleus accumbens. The dorsal striatum has been associated with reward-based action selection^16^ — and is thus thought to play a key role in instrumental learning tasks, such as ours, by maintaining information about action-contingent response-reward associations to guide future choices based on the outcomes of past ones — whereas the ventral portion of the striatum, the nucleus accumbens, has been more implicated in classical conditioning^16^. By blunting prediction error signals in the dorsal striatum, acute stress may thus impair learning of stimulus-response-reward associations, which are crucial to perform our reinforcement-learning task; yet, it is possible that acute stress affects distinct regions of the striatum, and the functions they support, differently^36,37^. Further studies are required to better understand the impact of acute stress on the neurocomputational mechanisms of classical and instrumental reward and punishment learning and could focus not only on the striatum, but also on other brain areas, such as the anterior insula or habenula, which are known to play a key role in punishment avoidance^15,47,54^.

In this study, we induced acute stress in participants, using a repetitive and uncontrollable sound, whilst they completed a reinforcement-learning task. Such stressor was previously validated outside the scanner using self-reported stress levels and concomitant skin conductance responses [for a thorough discussion about the choice and validation of the stressor see Carvalheiro et al., (2021)^7^]. In the current study, the stressor increased self-reported stress levels, although to a lesser extent than outside the scanner. One potential explanation for this discrepancy is that, inside the scanner, the stressor was not as salient as it was outside the scanner. We used additional visual cues as warning signals and coloured backgrounds, in an attempt to amplify the effects of stress manipulation. Importantly, the manipulation seemed to be particularly effective in a proportion of participants, based on their self-reported stress levels. In this study we used a self-report measure to capture the ‘subjective state of sensing potentially adverse changes in the environment’^26^. Although we cannot assume that the same processes governing physiological stress explain subjective feelings of stress^55,56^, the subjective emotional experience of stress plays an important role in the stress response^57^. Due to technical limitations, we did not analyse physiological responses in this study [but see Carvalheiro et al., (2021)^7^ for a skin conductance measure obtained for the same stress manipulation]; still, inclusion of such variables in future research could be valuable.

To avoid the potential confounding effects of menstrual-cycle-dependent variation on stress responsivity^58^ as well as on reward and punishment learning^59,60^, only men were included in this study. Our behavioural findings seem to be in line with previous reports showing that acute stress disrupts reward learning in women^1–3,5,6^, but further studies are needed to assess whether acute stress affects the same neurocomputational mechanisms of reinforcement learning in both men and women. Furthermore, given that our stress manipulation did not increase stress levels in all participants, future studies should explicitly account for individual differences, and for the modulatory role of those individual differences on the neural mechanisms that underlie reinforcement learning under acute stress.

In sum, we present evidence that acute stress blunts prediction error signals in the dorsal striatum during reinforcement learning. This effect seems to be mostly driven by decreased positive prediction error signals, which might explain why individuals learn worse from the rewarding outcomes of their choices when under acute stress. Our findings can thus contribute to a better understanding of the neural mechanisms that underlie the deleterious impact of acute stress on reward learning. Ultimately, this study may offer important mechanistic insights into the impact of acute stress in everyday life as well as on designing appropriate interventions^61,62^.

## Methods

### Participants

We scanned a total of 42 right-handed male participants with no reported history of neurological or psychiatric disorders. One participant was excluded due to incidental findings and 4 participants were excluded due to technical problems during the scanning session. We assessed whether the stress manipulation increased stress levels by comparing the self-reported stress levels of the remaining 37 participants that completed the task in the stress and control conditions. Self-reported stress levels were higher in the stress condition than in the control condition in 23 participants. Thus, we report results from data analyses of these 23 participants (age range = 18 – 29 years; *M* = 23.0 years, *SD* = 3.3 years). For completeness, we also analysed the data from the total sample (n = 37); those analyses can be found in the Supplementary Information.

All participants provided their informed consent before the experimental session. All experimental procedures were approved by the Ethics Committee of Hospital of Braga.

### Reinforcement-learning task

After a short practice (12 trials) outside the scanner to familiarise participants with the task timings and response keys, participants completed four blocks of an adapted version of a well-established reinforcement-learning task^15^ whilst inside the scanner (Fig. 1). The task was divided in two runs, each run consisting of a stress block and a control block. Stress and control blocks were administered alternately and in a counterbalanced order across the two runs. Each block included three pairs of abstract stimuli, and each pair of stimuli was presented 24 times, totalling 72 trials per block. New abstract stimuli were used in each block. Each pair of stimuli was associated with a valence: one pair of stimuli was associated with gains (gain 0.5€ or no change), a second pair was associated with losses (loss 0.5€ or no change), and a third pair was associated with neutral, or non-financial outcomes (look at a 0.5€ coin or no change). The outcome probabilities were reciprocally 0.75 and 0.25 for the stimuli in each of the three pairs. On each trial, one pair was randomly presented on the MRI screen, with one stimulus from the pair on the left and the other on the right of a central fixation cross (the stimuli position was counterbalanced across trials). Participants were instructed to choose between the two visual stimuli displayed on the screen to maximize payoffs. Missing choices occurred when participants did not press the response keys within 2000 ms (total of 0.20% missing choices: 8 in the stress condition and 5 in the control condition, in a total of 6624 trials) and were signaled with a “Missed” message (no other outcome was provided). Missing choices were not considered for behavioural data analyses. Before starting the task, participants were informed that they would be paid the amount of money obtained during their most profitable block, although they all left with the same fixed compensation (15€) for their participation. The experiment was programmed and presented with Cogent 2000 (http://www.vislab.ucl.ac.uk/cogent.php) implemented in MATLAB R2015a (MathWorks).

### Acute-stress manipulation

During the scanning session, participants performed two blocks of the reinforcement-learning task whilst exposed to a stressor (i.e., stress condition) and two blocks without the stressor (i.e., control condition) (Fig. 1). By exposing participants to the stressor during the task, we aimed to make sure that acute stress was contingent on the learning processes. To elicit stress responses, we exposed participants to a predictable, but uncontrollable auditory stimulus: a constant alarm (“Annoying modern office building alarm.wav”, retrieved from freesound.org, and programmed to loop uninterruptedly), played through the scanner with the volume set to the maximum. This uncontrollable sound was always constant and repetitive, to minimise the potential entanglement between stress and distraction, as evidence suggests that unpredictable changes in sound sequences seem to induce distraction more robustly^63–65^. Stress blocks were further signalled by a warning sign and a red background (Fig. 1a), and control blocks were signalled by a safe sign and blue background (Fig. 1b).

Stress levels were assessed by asking participants at the end of each block to rate how stressed they felt during that block on a scale of 1 (nothing) to 9 (extremely). We showed in a previous behavioural study that this stress manipulation increased self-reported stress levels and skin conductance responses rate in men^7^.

### Task performance analyses

To examine the impact of acute stress on behavioural choice performance during the reinforcement-learning task, we applied a generalized linear mixed-effects (glme) model to participants’ trial-by-trial choice data (with correct and incorrect choices coded as 1 and 0, respectively). We used a “logit” link function to account for the binomial distribution of the data. We included as predictor variables in the glme model: condition (stress or control), valence (gain or loss), block number (1 or 2), trial number (1 to 24), and the interaction of interest (condition × valence). The glme included a fixed intercept, as well as random intercepts for each participant. We fitted the glme model to the behavioural data using MATLAB’s *fitglme* function and performed planned post-hoc analyses via contrast matrices using MATLAB’s *coefTest* function.

### fMRI data acquisition and preprocessing

A Siemens Verio 3T MRI scanner at the Clinical Academic Center – Braga with a 32-channel head coil was used to acquire a 5.5 min 3D T1-weighted anatomical scan and multislice T2*-weighted echo planar images (EPIs) with BOLD contrast. The T2* EPI sequence used the following acquisition parameters: field of view = 200 x 200 mm, matrix size = 66 x 66 mm, interleaved slice order acquisition, 42 slices with slice thickness of 3 mm with no gap between slices, flip angle of 60°, echo time of 22 ms, and repetition time of 2000 ms. Functional task-related data were acquired in two runs, separated by a short break during which participants remained inside the scanner in the same position. Fieldmaps were acquired for use in the unwarping stage of data preprocessing. Imaging data were analysed using SPM12 (www.fil.ion.ucl.ac.uk/spm). Data preprocessing followed a standard sequence: the first five volumes were discarded, and data were realigned to the sixth volume, unwarped using a fieldmap (normalized to the Montreal Neurological Institute, MNI, template), and coregistered to the participant’s own anatomical image. The anatomical images were normalized using a unified segmentation procedure^66^, combining segmentation, bias correction, and calculation of the wrapping or distortions needed to map the anatomical image into Montreal Neurological Institute space (i.e., deformation fields), and then applying these warps to the EPI data. The voxel size was resampled to 1.5 × 1.5 × 1.5 mm. Last, a Gaussian kernel of 8 mm FWHM was applied to smooth the images spatially.

### fMRI data analyses

#### Primary general linear model

The primary fMRI analyses were based on a single general linear model, as in previous studies that used a similar reinforcement-learning task^15,37,67^. Each trial was modelled as having two time points: stimuli and outcome onsets. Note that, although our analyses focused on the prediction errors at the onset of outcomes, the onsets of stimuli were also modelled, to account for likely shared variance between BOLD signals at the time of the stimuli and outcomes. Separate regressors were created for the 6 types of stimuli [2 conditions (stress/control) × 3 valences (gain/loss/neutral)] and the 6 types of outcomes [2 conditions (stress/control) × 3 valences (gain/loss/neutral)] in each run (see Supplementary Fig. 5 for an example of a first-level design matrix); the regressors were modelled as stick functions and convolved with SPM’s canonical hemodynamic response function^15^. Each time point was regressed with a parametric modulator, separately for gain and loss trials: stimuli onset was modulated by the value of the chosen option, *Q*_*chosen*_(*t*); and, importantly, outcome onset was modulated by the prediction error, *δ*(*t*). Such values and predictions errors were estimated trial-wise using a well-established reinforcement-learning model^41^. Briefly, in this reinforcement-learning model, the value of the chosen stimulus, *Q*_*chosen*_, is updated on each trial, *t*, according to the following learning rule: *Q*_*chosen*_(*t* + 1) = *Q*_*chosen*_(*t*) + α \**δ*(*t*). The prediction error, *δ* (*t*), is the difference between the actual and the expected outcome: *δ*(*t*) = *r*(*t*) −*Q*_*chosen*_(*t*), where the reinforcement *r*(*t*) is either 0.5, 0, or -0.5. The used reinforcement-learning model included separate learning rates for positive (*α*^*+*^) and negative (*α*^*-*^) prediction errors to account for the differential signalling of positive and negative prediction errors^13,14^. The reinforcement-learning model also included the inverse temperature parameter, *β*, which accounts for randomness in choice selection (see Supplementary Information for a detailed description of the reinforcement-learning model). Values and prediction errors were estimated using the parameters *α*^±^ and *β* estimated for each subject in each condition and used as separate parametric modulators of neural activity at the time of stimuli and outcomes, respectively, either in gain or loss trials, in each condition. We also included an additional regressor to model missed trials, when participants did not select one of the two symbols and there was no outcome. For participants with visible headmotion in a particular scan (scans with >1 mm or 1° movement relative to the next) an extra regressor was included. Those images were removed and replaced with an image created by interpolating the two adjacent images to prevent distortion of the between-subjects mask (seven participants with visible headmotion; less than 1% of the total time series for each of them). Six headmotion parameters modelled the residual effects of headmotion. Data were high-pass filtered at 128s to remove low-frequency drifts, and the statistical model included an AR(1) autoregressive function to account for autocorrelations intrinsic to the fMRI time series.

Our primary analyses focused on prediction errors at outcome. First-level contrast images were calculated by applying appropriate linear contrasts to the parametric modulators of interest — prediction errors — and were entered into second-level analyses. Second-level one-sample *t*-tests were conducted for each contrast using the summary-statistics approach to random-effects analysis. Regions of interest (ROI) analyses in the dorsal striatum and nucleus accumbens were conducted using an initial threshold of *p* < 0.001 (uncorrected) and responses were considered significant if they survived voxel-level small-volume family-wise error correction (SVC-FWE) at *p* < 0.05. The *a priori* ROIs — dorsal striatum and nucleus accumbens — were anatomically defined using masks. Specifically, a bilateral mask for the dorsal striatum was defined using a conjunction of the left and right putamen and caudate from the automated anatomical labelling (AAL) atlas. A bilateral mask for the nucleus accumbens was defined using a conjunction of the left and right nucleus accumbens from the Individual Brain Atlases using Statistical Parametric Mapping (IBASPM). As the bilateral nucleus accumbens mask had a slight overlap with the dorsal striatum mask, we subtracted the mask of the nucleus accumbens from the dorsal striatum mask. The atlases and the conjunctions were implemented using the WFU PickAtlas Toolbox in SPM12. Individual BOLD estimates (i.e., regression slopes) of prediction error parametric modulators were extracted from significantly activated clusters using the MarsBaR toolbox^68^.

#### Subsidiary general linear models

To better understand the impact of acute stress on prediction error signals, we generated two subsidiary general linear models. Note that these two subsidiary models did not include any parametric modulators, as our purpose was to visualise how the BOLD response varied along different magnitudes of prediction errors.

For the first subsidiary model, we split prediction errors into four equally sized bins. The boundaries of the bins did not differ significantly between the stress and control conditions, in gain (all *p* > 0.071, paired *t*-tests) nor in loss (all *p* > 0.13, paired *t*-tests) trials (Supplementary Table 2). Specifically, this first subsidiary general linear model included separate regressors for trials corresponding to each bin, in each valence (gain and loss) and condition (stress and control), modelled at the stimuli and outcome onset (as in the primary general linear model) resulting in thirty-two regressors, plus regressors for neutral trials in each condition, missing trials (if applicable) and headmotion (and visible headmotion, if applicable), for each run.

For the second subsidiary model, we split prediction errors into negative and positive. This model included separate regressors for trials corresponding to negative and positive prediction errors, in each valence (gain and loss) and condition (stress and control), modelled at the stimuli and outcome onset (as in the primary general linear model) resulting in sixteen regressors, plus regressors for neutral trials in each condition, missing trials (if applicable) and headmotion (and visible headmotion, if applicable), for each run.

In both subsidiary models, the average BOLD estimates at the outcome onset (when prediction errors occur) were extracted from the significant dorsal striatum cluster identified in the primary general linear model, using the MarsBaR toolbox^68^.

## Supporting information

Supplementary Information_stress_PEs

## Acknowledgments

This work was supported by grants from the Portuguese Foundation for Science and Technology (FCT) to ASC [SFRH/BPD/94970/2013, PTDC/MHC-PCN/2296/2014, co-financed by FEDER through COMPETE2020 under the PT2020 Partnership Agreement (POCI-01-0145-FEDER-016747)] and to AM (IF/00750/2015). JC was supported by a FCT Ph.D. fellowship (PD/BD/128467/2017). This study was conducted at the Psychology Research Centre (PSI/01662), School of Psychology, University of Minho, supported by the Foundation for Science and Technology (FCT) through the Portuguese State Budget (UID/PSI/01662/2020).

## Author Contributions

J.C., A.M., and A.S.C. designed the experiment. J.C. collected the data. J.C. analysed the data. J.C., V.C., A.M., and A.S.C. interpreted the data. J.C. drafted the manuscript, and all authors edited the manuscript.

## References

1. Bogdan, R. & Pizzagalli, D. A. Acute stress reduces reward responsiveness: implications for depression. Biological Psychiatry 60, 1147–1154 (2006).

2. Bogdan, R., Santesso, D. L., Fagerness, J., Perlis, R. H. & Pizzagalli, D. A. Corticotropin-releasing hormone receptor type 1 (CRHR1) genetic variation and stress interact to influence reward learning. The Journal of Neuroscience 31, 13246–13254 (2011).

3. Berghorst, L. H., Bogdan, R., Frank, M. J. & Pizzagalli, D. A. Acute stress selectively reduces reward sensitivity. Frontiers in Human Neuroscience 7, (2013).

4. Ehlers, M. R. & Todd, R. M. Acute psychophysiological stress impairs human associative learning. Neurobiology of Learning and Memory 145, 84–93 (2017).

5. Morris, B. H. & Rottenberg, J. Heightened reward learning under stress in generalized anxiety disorder: A predictor of depression resistance? Journal of Abnormal Psychology 124, 115–127 (2015).

6. Paret, C. & Bublatzky, F. Threat rapidly disrupts reward reversal learning. Behaviour Research and Therapy 131, 103636 (2020).

7. Carvalheiro, J., Conceição, V. A., Mesquita, A. & Seara-Cardoso, A. Acute stress impairs reward learning in men. Brain and Cognition 147, 105657 (2021).

8. Cremer, A., Kalbe, F., Gläscher, J. & Schwabe, L. Stress reduces both model-based and model-free neural computations during flexible learning. NeuroImage 117747 (2021).

9. de Berker, A. O. et al. Acute stress selectively impairs learning to act. Scientific Reports 6, 29816 (2016).

10. Petzold, A., Plessow, F., Goschke, T. & Kirschbaum, C. Stress reduces use of negative feedback in a feedback-based learning task. Behavioral Neuroscience 124, 248–255 (2010).

11. O’Doherty, J. P., Hampton, A. & Kim, H. Model-based fMRI and its application to reward learning and decision making. Annals of the New York Academy of Sciences 1104, 35–53 (2007).

12. Sutton, R. & Barto, A. Reinforcement Learning: An Introduction. (MIT Press, 1998).

13. Maia, T. V. & Frank, M. J. From reinforcement learning models to psychiatric and neurological disorders. Nature Neuroscience 14, 154–162 (2011).

14. Schultz, W., Dayan, P. & Montague, P. R. A neural substrate of prediction and reward. Science 275, 1593–9 (1997).

15. Pessiglione, M., Seymour, B., Flandin, G., Dolan, R. J. & Frith, C. D. Dopamine-dependent prediction errors underpin reward-seeking behaviour in humans. Nature 442, 1042–1045 (2006).

16. O’Doherty, J. et al. Dissociable roles of ventral and dorsal striatum in instrumental conditioning. Science 304, 452–454 (2004).

17. Valentin, V. V. & O’Doherty, J. P. Overlapping prediction errors in dorsal striatum during instrumental learning with juice and money reward in the human brain. Journal of Neurophysiology 102, 3384–3391 (2009).

18. Schönberg, T., Daw, N. D., Joel, D. & O’Doherty, J. P. Reinforcement learning signals in the human striatum distinguish learners from nonlearners during reward-based decision making. The Journal of Neuroscience 27, 12860–12867 (2007).

19. Daw, N. D. & Tobler, P. N. Value learning through reinforcement: The basics of dopamine and reinforcement learning. in Neuroeconomics: Decision Making and the Brain (eds. Glimcher, P. W. & Fehr, E.) 283–298 (Elsevier Inc., 2014).

20. Glimcher, P. W. Understanding dopamine and reinforcement learning: The dopamine reward prediction error hypothesis. Proceedings of the National Academy of Sciences 108, 15647 (2011).

21. Pruessner, J. C., Champagne, F., Meaney, M. J. & Dagher, A. Dopamine release in response to a psychological stress in humans and its relationship to early life maternal care: a positron emission tomography study using [11C]raclopride. The Journal of Neuroscience 24, 2825–2831 (2004).

22. Booij, L. et al. Dopamine cross-sensitization between psychostimulant drugs and stress in healthy male volunteers. Translational Psychiatry 6, e740 (2016).

23. Cabib, S. & Puglisi-Allegra, S. The mesoaccumbens dopamine in coping with stress. Neuroscience and Biobehavioral Reviews 36, 79–89 (2012).

24. de Kloet, E. R., Joëls, M. & Holsboer, F. Stress and the brain: from adaptation to disease. Nature Reviews Neuroscience 6, 463–475 (2005).

25. Hermans, E. J., Henckens, M. J. A. G., Joëls, M. & Fernández, G. Dynamic adaptation of large-scale brain networks in response to acute stressors. Trends in Neurosciences 37, 304–314 (2014).

26. Joëls, M. & Baram, T. Z. The neuro-symphony of stress. Nature reviews. Neuroscience 10, 459–466 (2009).

27. Abercrombie, E. D., Keefe, K. A., DiFrischia, D. S. & Zigmond, M. J. Differential effect of stress on in vivo dopamine release in striatum, nucleus accumbens, and medial frontal cortex. Journal of Neurochemistry 52, 1655–1658 (1989).

28. Vaessen, T., Hernaus, D., Myin-Germeys, I. & van Amelsvoort, T. The dopaminergic response to acute stress in health and psychopathology: A systematic review. Neuroscience and Biobehavioral Reviews 56, 241–251 (2015).

29. Valenti, O., Lodge, D. J. & Grace, A. A. Aversive stimuli alter ventral tegmental area dopamine neuron activity via a common action in the ventral hippocampus. The Journal of Neuroscience 31, 4280–4289 (2011).

30. Anstrom, K. K. & Woodward, D. J. Restraint increases dopaminergic burst firing in awake rats. Neuropsychopharmacology 30, 1832–1840 (2005).

31. Anstrom, K. K., Miczek, K. A. & Budygin, E. A. Increased phasic dopamine signaling in the mesolimbic pathway during social defeat in rats. Neuroscience 161, 3–12 (2009).

32. Maia, T. V. & Frank, M. J. An integrative perspective on the role of dopamine in schizophrenia. Biological Psychiatry 81, 52–66 (2017).

33. Werlen, E. et al. Amphetamine disrupts haemodynamic correlates of prediction errors in nucleus accumbens and orbitofrontal cortex. Neuropsychopharmacology 45, 793–803 (2020).

34. Daberkow, D. P. et al. Amphetamine paradoxically augments exocytotic dopamine release and phasic dopamine signals. The Journal of Neuroscience 33, 452–463 (2013).

35. Porcelli, A. J. & Delgado, M. R. Stress and decision making: effects on valuation, learning, and risk-taking. Current Opinion in Behavioral Sciences 14, 33–39 (2017).

36. Robinson, O. J., Overstreet, C., Charney, D. R., Vytal, K. & Grillon, C. Stress increases aversive prediction error signal in the ventral striatum. Proceedings of the National Academy of Sciences 110, 4129–4133 (2013).

37. Treadway, M. T. et al. Association between interleukin-6 and striatal prediction-error signals following acute stress in healthy female participants. Biological Psychiatry 82, 570–577 (2017).

38. Palminteri, S. & Pessiglione, M. Opponent brain systems for reward and punishment learning: Causal evidence from drug and lesion studies in humans. in Decision Neuroscience: An Integrative Perspective 291–303 (Elsevier Inc., 2017).

39. Watabe-Uchida, M., Eshel, N. & Uchida, N. Neural circuitry of reward prediction error. Annual Review of Neuroscience 40, 373–394 (2017).

40. Oleson, E. B., Gentry, R. N., Chioma, V. C. & Cheer, J. F. Subsecond dopamine release in the nucleus accumbens predicts conditioned punishment and its successful avoidance. The Journal of Neuroscience 32, 14804 (2012).

41. Frank, M. J., Moustafa, A. A., Haughey, H. M., Curran, T. & Hutchison, K. E. Genetic triple dissociation reveals multiple roles for dopamine in reinforcement learning. Proceedings of the National Academy of Sciences 104, 16311–16316 (2007).

42. Diederen, K. M. J. et al. Dopamine modulates adaptive prediction error coding in the human midbrain and striatum. The Journal of Neuroscience 37, 1708–1720 (2017).

43. Doll, B. B., Hutchison, K. E. & Frank, M. J. Dopaminergic genes predict individual differences in susceptibility to confirmation bias. The Journal of Neuroscience 31, 6188–6198 (2011).

44. Frank, M. J. & Fossella, J. A. Neurogenetics and pharmacology of learning, motivation, and cognition. Neuropsychopharmacology 36, 133–152 (2011).

45. Grogan, J. P. et al. Effects of dopamine on reinforcement learning and consolidation in Parkinson’s disease. Elife 6, (2017).

46. Rutledge, R. B. et al. Dopaminergic drugs modulate learning rates and perseveration in Parkinson’s patients in a dynamic foraging task. The Journal of Neuroscience 29, 15104–15114 (2009).

47. Garrison, J., Erdeniz, B. & Done, J. Prediction error in reinforcement learning: a meta-analysis of neuroimaging studies. Neuroscience and Biobehavioral reviews 37, 1297–1310 (2013).

48. Sulzer, D., Cragg, S. J. & Rice, M. E. Striatal dopamine neurotransmission: Regulation of release and uptake. Basal Ganglia 6, 123–148 (2016).

49. Wightman, R. M. et al. Dopamine release is heterogeneous within microenvironments of the rat nucleus accumbens. The European Journal of Neuroscience 26, 2046–2054 (2007).

50. Grace, A. A. Dysregulation of the dopamine system in the pathophysiology of schizophrenia and depression. Nature Reviews Neuroscience 17, 524–532 (2016).

51. Bayer, H. M. & Glimcher, P. W. Midbrain dopamine neurons encode a quantitative reward prediction error signal. Neuron 47, 129–141 (2005).

52. Boureau, Y.-L. & Dayan, P. Opponency revisited: Competition and cooperation between dopamine and serotonin. Neuropsychopharmacology 36, 74–97 (2011).

53. Eisenegger, C. et al. Role of dopamine D2 receptors in human reinforcement learning. Neuropsychopharmacology 39, 2366–2375 (2014).

54. Salas, R., Baldwin, P., de Biasi, M. & Montague, P. R. BOLD responses to negative reward prediction errors in human habenula. Frontiers in Human Neuroscience 4, 36 (2010).

55. Campbell, J. & Ehlert, U. Acute psychosocial stress: does the emotional stress response correspond with physiological responses? Psychoneuroendocrinology 37, 1111–1134 (2012).

56. Ali, N., Nitschke, J. P., Cooperman, C. & Pruessner, J. C. Suppressing the endocrine and autonomic stress systems does not impact the emotional stress experience after psychosocial stress. Psychoneuroendocrinology 78, 125–130 (2017).

57. Birk, R. H. On stress and subjectivity. Theory & Psychology (2020).

58. Ossewaarde, L. et al. Neural mechanisms underlying changes in stress-sensitivity across the menstrual cycle. Psychoneuroendocrinology 35, 47–55 (2010).

59. Diekhof, E. K. & Ratnayake, M. Menstrual cycle phase modulates reward sensitivity and performance monitoring in young women: Preliminary fMRI evidence. Neuropsychologia 84, 70–80 (2016).

60. Dreher, J.-C. et al. Menstrual cycle phase modulates reward-related neural function in women. Proceedings of the National Academy of Sciences 104, 2465–2470 (2007).

61. Nair, A., Rutledge, R. B. & Mason, L. Under the hood: Using computational psychiatry to make psychological therapies more mechanism-focused. Frontiers in Psychiatry 11, 140 (2020).

62. Queirazza, F., Fouragnan, E., Steele, J. D., Cavanagh, J. & Philiastides, M. G. Neural correlates of weighted reward prediction error during reinforcement learning classify response to cognitive behavioral therapy in depression. Science Advances 5, eaav4962 (2019).

63. Hughes, R. W. Auditory distraction: A duplex-mechanism account. PsyCh Journal 3, 30–41 (2014).

64. Parmentier, F. B. R. The cognitive determinants of behavioral distraction by deviant auditory stimuli: a review. Psychological Research 78, 321–338 (2014).

65. Parmentier, F. B. R., Elford, G., Escera, C., Andrés, P. & Miguel, I. S. The cognitive locus of distraction by acoustic novelty in the cross-modal oddball task. Cognition 106, 408–432 (2008).

66. Ashburner, J. & Friston, K. J. Unified segmentation. Neuroimage 26, 839–851 (2005).

67. Kumar, P. et al. Impaired reward prediction error encoding and striatal-midbrain connectivity in depression. Neuropsychopharmacology 43, 1581–1588 (2018).

68. Brett M., Anton J.L., Valabregue R., & Poline J.B. Region of interest analysis using an SPM toolbox. 8th International Conference on Functional Mapping of the Human Brain. Available on CD-ROM. in NeuroImage (2002).

