## Supplementary Information_stress_PEs for "Acute stress blunts prediction error signals in the dorsal striatum during reinforcement learning"

**This PDF file includes:**

**Analyses of the total sample**

Manipulation check (Supplementary Fig. 1)

Task-performance analyses (Supplementary Fig. 2)

fMRI analyses (Supplementary Fig. 3 and Supplementary Fig. 4)

**Primary general linear model** (Supplementary Fig. 5)

**Computational modelling**

Reinforcement-learning model

Reinforcement-learning modelling results (Supplementary Fig. 6)

**Supplementary tables** (Supplementary Table 1 and Supplementary Table 2)

**Supplementary references**

### Analyses of the total sample

To check the success of the acute-stress manipulation, we collected self-reported stress levels at the end of each task block. As we were interested in examining the impact of acute stress on reinforcement learning, participants who reported to be non-responsive to the stress manipulation (i.e., who did not report higher stress levels in the stress condition than in the control condition) were excluded from the behavioural and neuroimaging analyses presented in the main text. For completeness, here we present data on self-reported stress levels (c.f. section ‘Manipulation check’), as well as on task-performance (c.f. section ‘Task-performance analyses’) and neuroimaging (c.f. section ‘fMRI analyses’) analyses, in the total sample.

#### Manipulation check

To assess whether the acute-stress manipulation increased stress levels in the total sample (n = 37), we conducted an analysis of variance (ANOVA) with condition (stress and control) and block (1 and 2) as within-subject factors. Self-reported stress levels differed significantly between conditions ($F_{1,36} = 4.53$, $p = 0.04$, $\eta_2 = 0.11$) (Supplementary Fig. 1), and there was no main effect of block ($F_{1,36} = 0.25$, $p = 0.62$, $\eta_2 = 0.007$). The condition × block interaction was also non-significant ($F_{1,36} = 2.63$, $p = 0.11$, $\eta_2 = 0.068$), suggesting that self-reported stress levels remained stable across blocks within each condition.

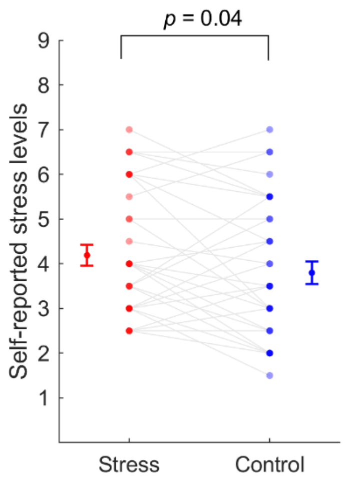


**Supplementary Fig. 1.** Self-reported stress levels in the stress (red) and control (blue) conditions in the total sample (n = 37). Connected dots represent data points from the same participant, and more transparent (opaque) dots represent less (more) overlapping data points; error bars displayed on the sides of those scatter plots indicate the mean ± standard error of the mean.

### Task-performance analyses

We conducted task-performance analyses on the trial-by-trial choice data from the total sample, using a generalized linear mixed-effects model, as described in the main manuscript. We found a significant condition × valence interaction ($\beta$ = -0.40, $p <$ 0.001, 95% CI = [-0.62, -0.18]) (Supplementary Fig. 2a). Planned post-hoc analyses revealed that under stress, comparatively to the control condition, participants performed worse when learning to obtain gains ($F_{1,\,7078}$= 7.47, $p$ = 0.0063), but better when learning to avoid losses ($F_{1,\,7078}$ = 6.02, $p$ = 0.014) (Supplementary Fig. 2b). These somewhat unexpected results concerning punishment learning were driven by the inclusion (in the total sample) of participants who reported lower levels of stress in the stress condition than in the control condition (n = 11) and who performed better, both in gain ($F_{1,\,2097}$ = 4.63, $p$ = 0.031) and loss ($F_{1,\,2097}$ = 25.97, $p$ < 0.001) trials, in the stress condition than in the control condition — note that, like in our subsample of interest, those participants had a worse performance in the condition in which they reported higher stress levels.

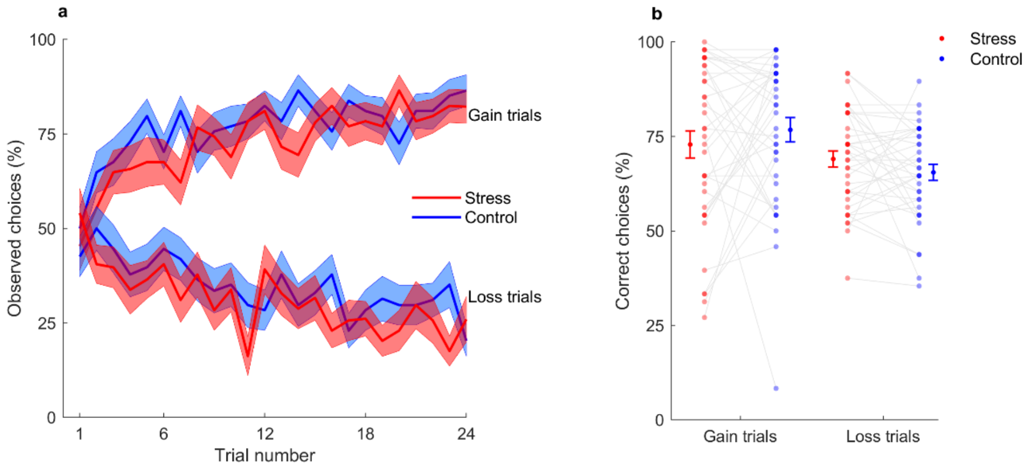


**Supplementary Fig. 2.** Task performance in the total sample. **(a)** Learning curves represent the trial-by-trial percentage of participants (n = 37) who chose the correct gain stimulus (associated with a probability of 0.75 of winning 0.5€; upper part of the graph) and the incorrect loss stimulus (associated with a probability of 0.75 of losing 0.5€; lower part of the graph), in the stress (red) and control (blue) conditions. Each central line represents the mean and the filled area the ± standard error of the mean. **(b)** Percentage of correct gain and loss choices per participant (n = 37) in stress and control conditions (averaged across blocks). Connected dots represent data points from the same participant, and more transparent (opaque) dots represent less (more) overlapping data points; error bars displayed on the sides of those scatter plots indicate the mean ± standard error of the mean.

### fMRI analyses

We repeated the neuroimaging analyses, as described in the main manuscript, for the total sample. Specifically, we took the control > stress contrast to second-level analyses but now including data from the total sample (n = 37). We replicated our main finding that acute stress decreased the parametric modulation of prediction errors in the dorsal striatum ([$x$ = 32, $y$ = 0, $z$ = 12], $z$ = 4.83, $k$ =45, $p$ = 0.002, SVC-FWE) (Supplementary Fig. 3a). No significant responses were found in the nucleus accumbens for the control > stress contrast. The fMRI contrast that tested the condition (control and stress) × trial valence (gain and loss) interaction did not show any significant valence-dependent differences in the dorsal striatum nor in the nucleus accumbens.

To explore whether acute stress preferentially blunted positive or negative prediction errors, we conducted subsidiary analyses in the total sample, as described in the main text. Subsidiary data indicated that acute stress differentially affected positive and negative prediction error signals (condition × prediction error valence interaction, $F_{1,36} = 4.90$, $p = 0.033$, $\eta^2 = 0.12$), such that acute stress decreased positive prediction error signals, but not negative prediction error signals (Supplementary Fig. 3b).

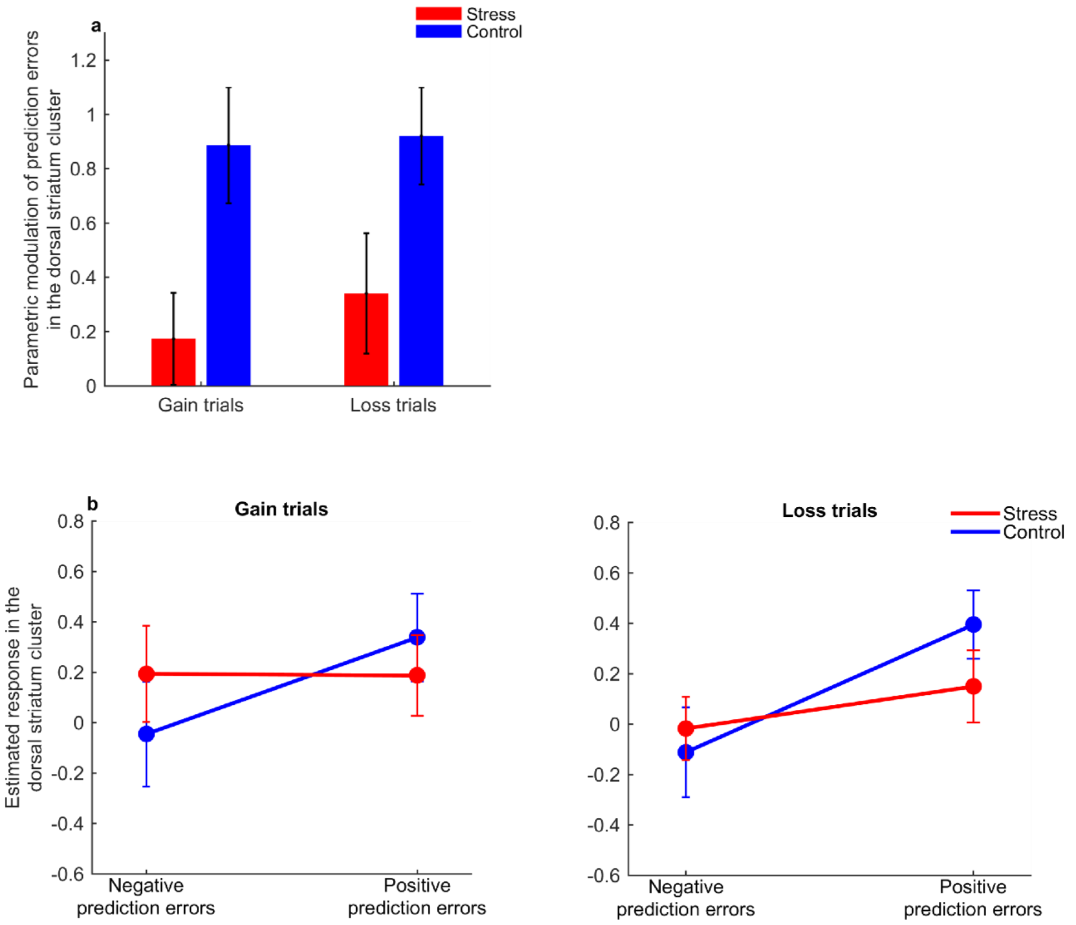


**Supplementary Fig. 3.** Effects of acute stress on prediction error signalling in the dorsal striatum in the total sample**. (a)** Bars depict the parameter estimates (i.e., regression slopes) for the BOLD response at the dorsal striatum cluster modulated by trial-by-trial prediction errors, in gain and loss trials, across the stress (red) and control (blue) conditions (n = 37). **(b)** Graphs illustrate the modulation of the BOLD response by prediction errors, during gain (left) and loss (right) trials, at the dorsal striatum cluster in the stress (red) and control (blue) conditions in the total sample (n = 37). Prediction errors were divided into negative and positive, and the BOLD response estimates at the outcome onset were extracted from the dorsal striatum cluster identified in the primary parametric modulation model for each participant. Error bars indicate the mean ± standard error of the mean.

Additionally, as the total sample of participants was characterized by variability in stress responses, with participants reporting to be less or more stressed in the stress condition relative to the control condition, we conducted exploratory analyses to probe whether the impact of stress on prediction errors signalling was associated with individual differences in stress levels across the total sample. For this, we extracted the parameter estimates (i.e., regression slopes) for the modulation of the BOLD response by prediction errors, in the dorsal striatal cluster identified in the above parametric modulation analysis, and correlated those parameter estimates with the difference in self-reported stress levels between the stress and control conditions. We found that prediction error signals during gain trials in the stress condition correlated negatively with the difference between self-reported stress levels in the stress and control condition (Pearson's correlation: $r$= -0.55, $p < 0.001$; Spearman's correlation: $rs$= -0.52, $p$ = 0.001) (Supplementary Fig. 4).

This tentative finding suggests that participants who reported the greatest increase in stress levels in response to the acute stressor, relative to control condition, also showed the greatest reduction in signalling of prediction errors in the dorsal striatum during reward learning when under acute stress.

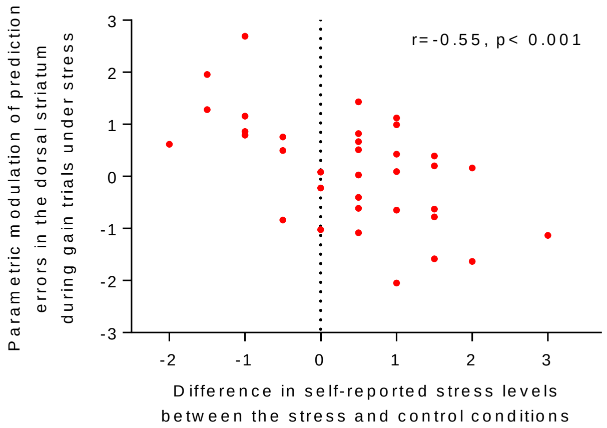


**Supplementary Fig. 4.** Scatterplot of the difference between self-reported stress levels in the stress and control condition and parameter estimates (i.e., regression slopes) of the BOLD response in the dorsal striatum cluster modulated by prediction errors during gain trials in the stress condition (n = 37). The dashed vertical line divides the plot into participants who reported to be responsive to the stress manipulation and participants who did not report to be responsive to the stress manipulation.

**Primary general linear model**

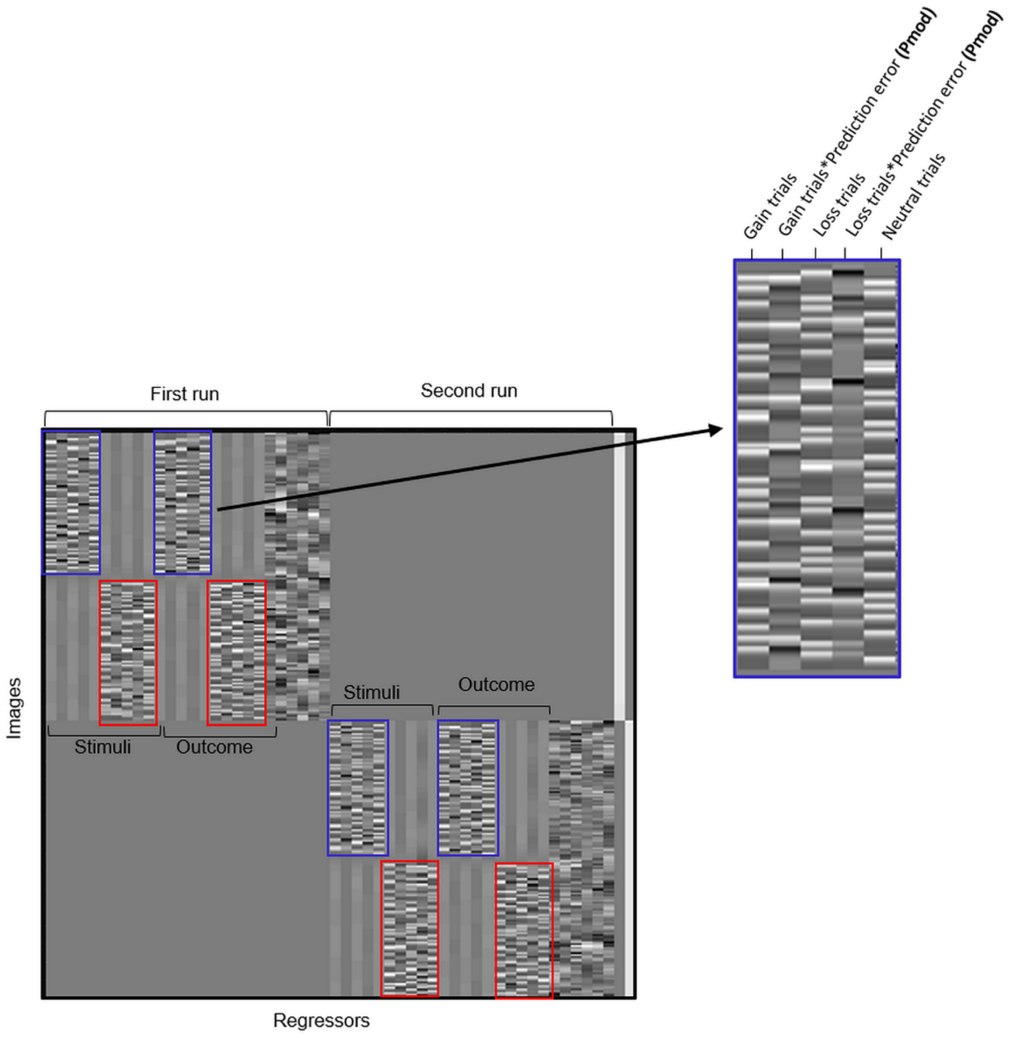


**Supplementary Fig. 5**. Example of a first-level SPM design matrix showing the regressors across the two runs of the task for an exemplary subject: columns (left to right) represent : 1) onsets of stimuli in gain and loss trials — which were parametrically modulated by chosen option values — and neutral trials, in the control (blue) and stress (red) conditions; 2) onsets of outcomes in gain and loss trials — which were parametrically modulated by prediction errors — and neutral trials, in the control (blue) and stress (red) conditions; 3) headmotion regressors (additional regressors for missing trials and/or visible headmotion were also included when needed; not depicted for this subject). The zoomed in regressors on the right are related to the onset of the outcomes in the control condition during the first run; the regressors of the parametric modulation (Pmod) by the prediction errors in gain and loss trials are depicted in the second and fourth columns, respectively.

### Computational modelling

#### Reinforcement-learning model

Prediction errors were calculated via a reinforcement-learning model that has been extensively used to investigate the behavioural and neural impact of pharmacological manipulations and genetic variations in the dopaminergic system in humans[1–6]. The used reinforcement-learning model included separate learning rates for positive ($\alpha^+$) and negative ($\alpha^-$) prediction errors, to account both for the differential firing of dopaminergic neurons for positive and negative prediction errors and the differential effects of dopamine onto the plasticity of the corticostriatal synapses implicated in action-value learning[7,8]. This model also included the inverse temperature parameter, $\beta$, which controls randomness in choice selection[9,10].

In the context of our task, the used reinforcement-learning model[4] assumes that each participant gradually learns the value of choosing a given stimulus (say A or B) from a given pair of stimuli (here, "gain" or "loss" stimuli pairs, as the "neutral" pair of stimuli always yielded null monetary outcomes) as a function of both the outcome that was obtained on that trial following stimulus selection and the participant's prior outcome expectation, as quantified by a $Q$-value. Specifically, each pair-stimulus value, or $Q$-value, was initialized to zero, and for each trial, $t$, within that pair of stimuli, the value of the chosen stimulus (say A was chosen) was updated according to:

$$Q_A\,(t+1) = \; Q_A(t) + \; \alpha * \delta(t),$$

where $\delta$ was the prediction error:

$$\delta(t) = r(t) - Q_A\,(t),$$

where $r(t)$ was 0.5 for winning 0.5€, 0 for getting nothing, and -0.5 for losing 0.5€. The learning rate, $\alpha$ , was given by:

$$\alpha = \begin{cases} \alpha^+, & if\ \delta(t) > 0 \\ \alpha^-, & if\ \delta(t) < 0 \end{cases},$$

where $\alpha^{+}$ and $\alpha^{-}$ were the learning rates for positive and negative prediction errors, respectively[4].

The probability of choosing one stimulus over another (say A over B) was given by the *softmax* equation:

$$P_A(t) = \frac{e^{[Q_A(t)*\beta]}}{e^{[Q_A(t)*\beta]} + e^{[Q_B(t)*\beta]}} ,$$

where the $\beta$ parameter, or inverse temperature, reflects the randomness of choices. The lower the $\beta$, the higher the randomness of choices.

We modelled participants' trial-by-trial behaviour in the stress and control conditions using this well-established, biologically inspired reinforcement-learning model[4]. Model fitting involved estimating the values of the parameters ($\alpha^{+}$, $\alpha^{-}$, and $\beta$) that best accounted for the respective trial-by-trial choices in each condition. We estimated the best-fitting model parameters ($\alpha^{+}$, $\alpha^{-}$, and $\beta$) for each subject in each condition using maximum *a posteriori* estimation[9]. Specifically, to optimize model parameters, we drew the learning rates from Beta distributions [*Beta* (1.1, 1.1)] and the inverse temperature from a Gamma distribution [*Gamma* (1.2, 5)][11,12]. We then used MATLAB's *fmincon* function, initialized at 100 random starting points of the parameter space, to search for the parameter values that minimized the negative log posterior of the observed sequence of choices, given the previously observed outcomes, with respect to different settings of the model parameters[9].

**Reinforcement-learning modelling results**

In this study, we were interested in examining the impact of acute stress on the neural coding of prediction errors. For completeness, we also assessed whether acute stress affected the parameters estimated by the reinforcement-learning model, as we had previously shown that acute stress decreased the learning rate for positive prediction errors using a similar version of the reinforcement-learning task[12]. Specifically, we conducted repeated-measures ANOVAs with condition (stress and control) and valence of the prediction error (positive and negative) as within-subject factors. We did not find significant differences in the learning rates between the stress and control conditions (main effect of condition: $F_{1,22} = 0.095$, $p = 0.76$, $\eta^2 = 0.004$). There was a main effect of valence, such that the learning rate for positive prediction errors was significantly higher than the learning rate for negative predictions errors in both conditions ($F_{1,22} = 12.26$, $p = 0.002$, $\eta^2 = 0.36$). The condition × valence interaction was non-significant ($F_{1,22} = 0.90$, $p = 0.35$, $\eta^2 = 0.039$). The inverse temperature parameter also did not differ between the stress and control conditions ($F_{1,22} = 2.24$, $p = 0.15$, $\eta^2 = 0.093$). We further analysed the products between each learning rate ($\alpha^{\pm}$) and the inverse temperature ($\beta$)., because in reinforcement-learning models $\alpha^{\pm}$ and $\beta$ tend to be inversely coupled[9], and, as a result, the parameters viewed separately can have larger estimation errors, while their product tends to be more reliably estimated, and thus better recovered[9,13,14]. We did not find evidence for significant differences in the product between learning rates and the inverse temperature parameter ($\alpha^{\pm} * \beta$) between the stress and control conditions (main effect of condition: $F_{1,22} = 3.02$, $p = 0.096$, $\eta^2 = 0.12$) and the condition × valence interaction was also non-significant ($F_{1,22} = 0.30$, $p = 0.59$, $\eta^2 = 0.014$). Importantly, the version of the reinforcement-learning task used in our previous study, where we found a decreased learning rate for positive prediction errors under acute stress[2], differed from the version of the task applied in the present study in some important methodological aspects, which contributes to explain the absence of significant results. Specifically, as in this study we were particularly interested in the early phase of learning, and we aimed at avoiding that the task being solved inside the scanner was too repetitive and long, we reduced the number of times we presented each pair of stimuli during each block from 40 to 24[15]. We also changed the outcome probabilities from 0.80/0.20 to 0.75/0.25[15] to elicit more non-negligible prediction errors, as in the present study we were specifically interested in examining the neural correlates of prediction errors. It is thus possible that by decreasing the number of trials and/or changing the outcome probabilities, we were not able to obtain parameter estimates as stable[14] as in our previous study[12]. In addition, our sample size (n = 23) was smaller than in our previous behavioural study (n = 62), which may have led to reduced power to detect significant effects of the stressor (which was itself perceived less intensely on this study; see Discussion on the main manuscript) on parameter estimates.

Although our study did not detect an effect of acute stress on the parameter estimates, we conducted two additional checks to validate the used reinforcement-learning model. First, we validated the used reinforcement-learning model by confirming that the probability of choices estimated under the reinforcement-learning model followed the same pattern of the actual observed choices. Specifically, we computed trial-by-trial choice probabilities for all participants using the best-fitting set of parameters in each condition (the actual observed choices and outcomes were used to update the choice probabilities). To assess whether the probabilities of choosing the correct answer estimated by the reinforcement-learning model (for each subject, the choice probabilities were averaged across gains or loss trials in each condition) followed the same pattern as the actual correct observed choices, we conducted Pearson's correlations for gains and losses in both conditions, using the respective mean percentages. Results confirmed that the choice probabilities estimated under the reinforcement-learning model showed a close correspondence with the actual observed choices across gain and loss trials in both conditions (gains in the stress condition: r =

0.99, $p < 0.0001$; gains in the control condition: r = 0.97, $p < 0.0001$; losses in the stress condition: r = 0.53, $p = 0.0087$; losses in the control condition: r = 0.64, $p = 0.0094$).

Then, we conducted parameter-recovery analyses. Specifically, we simulated the task-choice behaviour of 23 virtual participants using the parameter values that we had estimated for each of the 23 participants in each condition. We ran 100 simulations. Then, for each simulation, we fitted the model to the virtual participants' data to estimate new (recovered) parameters. Finally, we tested the correlations between the original and recovered parameters using Pearson's correlations. Parameter-recovery analyses demonstrated that the results of our model-fitting procedure were robust both in the stress and control conditions (all r > 0.55, all $p < 0.047$; Supplementary Fig. 6).

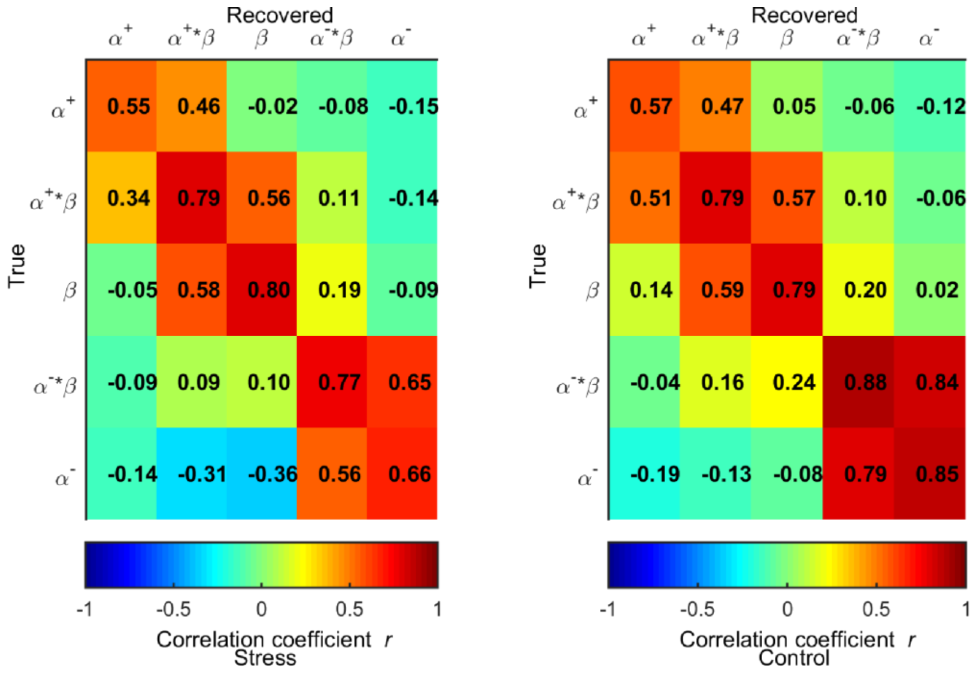


**Supplementary Fig. 6.** Correlation matrices between the estimated and recovered parameters averaged across 100 simulations in the stress (left) and control (right) conditions. The matrices include the values of Pearson's *r* for the correlations between the parameters that were used to generate the simulated data ("True") and those that were obtained by applying the parameter-estimation procedure to the simulated data ("Recovered").

### Supplementary tables

**Supplementary Table 1.** Positive parametric modulation of prediction errors. Regions are reported at a family-wise error (FWE) corrected $p < 0.05$ with a voxel-level whole-brain correction and a minimum of 10 contiguous voxels, unless otherwise stated. Spatial coordinates (x, y, z) are in the Montreal Neurological Institute (MNI) space, and regions were identified using the automated anatomical labelling (AAL) atlas. R = Right; L = Left. We also tested negative parametric modulations, but as we found no significant activations in our regions of interest, those results are not reported here.

[a] Regions reported at $p < 0.05$, using small-volume FWE voxel-level correction within bilateral masks of the referred regions (dorsal striatum or nucleus accumbens), following an initial threshold of p < .001 (uncorrected).

| Region | L/R | x | y | z | k | $p_{FWE}$ |
|---|---|---|---|---|---|---|
| **Gain trials in the control condition** | | | | | | |
| Occipital middle | R | 26 | -90 | 5 | 2809 | <.001 |
| Occipital inferior | L | -30 | -89 | 8 | 2419 | <.001 |
| Ventral striatum | L | -12 | 8 | -14 | 35 | .001 |
| Ventral striatum | R | 14 | 14 | -5 | 15 | .023 |
| Frontal superior | L | -20 | 32 | 53 | 36 | .001 |
| Fusiform | L | -32 | -36 | -23 | 11 | .046 |
| Dorsal striatum [a] | L | -11 | 9 | -11 | 1355 | <.001 |
| Dorsal striatum [a] | R | 17 | 12 | -6 | 791 | <.001 |
| Nucleus accumbens [a] | L | -12 | 8 | -14 | 114 | <.001 |
| Nucleus accumbens [a] | R | 14 | 12 | -8 | 135 | <.001 |
| **Gain trials in the stress condition** | | | | | | |
| Calcarine | R | 21 | -92 | 3 | 1092 | <.001 |
| Occipital middle | L | -15 | -98 | 0 | 736 | <.001 |
| Occipital inferior | R | 36 | -81 | -11 | 67 | <.001 |
| Occipital inferior | L | -32 | -84 | -12 | 14 | .015 |
| Fusiform | R | 30 | -42 | -20 | 23 | .003 |
| Caudate | L | -8 | 8 | -3 | 84 | <.001 |
| Dorsal striatum [a] | L | -8 | 8 | 3 | 850 | <.001 |
| Dorsal striatum [a] | R | 11 | 11 | -3 | 509 | .001 |
| Nucleus accumbens [a] | L | -11 | 6 | -8 | 95 | <.001 |
| Nucleus accumbens [a] | R | 8 | 8 | -8 | 119 | .001 |
| **Loss trials in the control condition** | | | | | | |
| Parietal Inferior | L | -44 | -32 | 42 | 21 | .005 |
| Dorsal striatum [a] | R | 27 | -8 | 12 | 629 | .007 |
| Dorsal striatum [a] | L | -18 | -12 | -11 | 199 | .014 |
| Dorsal striatum [a] | L | -27 | -11 | 12 | 297 | .022 |
| Dorsal striatum [a] | L | -18 | -2 | 24 | 65 | .035 |
| Nucleus accumbens [a] | L | -15 | 11 | -12 | 47 | <.001 |
| Nucleus accumbens [a] | R | 17 | 9 | -11 | 77 | .001 |
| **Loss trials in the stress condition** | | | | | | |
| Dorsal striatum [a] | L | -27 | -15 | 9 | 207 | .028 |

**Supplementary Table 2.** Median and boundaries of the magnitudes of the prediction errors for each of the four bins for each condition (stress or control) and each trial valence (gain or loss).

| | | **Median [lower bound; higher bound]** | | | |
|---|---|---|---|---|---|
| **Condition** | **Trial valence** | **Bin 1** | **Bin 2** | **Bin 3** | **Bin 4** |
| Stress | Gain | -0.39 [-0.43; -0.26] | -0.055 [-0.85; -0.018] | 0.080 [0.0037; 0.15] | 0.23 [0.17; 0.50] |
| | Loss | -0.46 [-0.50; -0.43] | -0.23 [-0.38; -0.12] | 0.01 [-0.062; 0.055] | 0.12 [0.088; 0.17] |
| Control | Gain | -0.40 [-0.44; -0.24] | -0.0078 [-0.036; 0.014] | 0.087 [0.049; 0.15] | 0.26 [0.19; 0.5] |
| | Loss | -0.56 [-0.5; -0.40] | -0.18 [-0.32; -0.12] | 0.017 [-0.059; 0.08] | 0.15 [0.097; 0.22] |

**Supplementary references**


1. Diederen, K. M. J. *et al.* Dopamine modulates adaptive prediction error coding in the human midbrain and striatum. *The Journal of Neuroscience* **37**, 1708–1720 (2017).
2. Doll, B. B., Hutchison, K. E. & Frank, M. J. Dopaminergic genes predict individual differences in susceptibility to confirmation bias. *The Journal of Neuroscience* **31**, 6188–6198 (2011).
3. Frank, M. J. & Fossella, J. A. Neurogenetics and pharmacology of learning, motivation, and cognition. *Neuropsychopharmacology* **36**, 133–152 (2011).
4. Frank, M. J., Moustafa, A. A., Haughey, H. M., Curran, T. & Hutchison, K. E. Genetic triple dissociation reveals multiple roles for dopamine in reinforcement learning. *Proceedings of the National Academy of Sciences* **104**, 16311–16316 (2007).
5. Grogan, J. P. *et al.* Effects of dopamine on reinforcement learning and consolidation in Parkinson's disease. *Elife* **6**, (2017).
6. Rutledge, R. B. *et al.* Dopaminergic drugs modulate learning rates and perseveration in Parkinson's patients in a dynamic foraging task. *The Journal of Neuroscience* **29**, 15104–15114 (2009).
7. Maia, T. V. & Frank, M. J. From reinforcement learning models to psychiatric and neurological disorders. *Nature Neuroscience* **14**, 154–162 (2011).
8. Frank, M. J. & O'Reilly, R. C. A mechanistic account of striatal dopamine function in human cognition: Psychopharmacological studies with cabergoline and haloperidol. *Behavioral Neuroscience* **120**, 497–517 (2006).
9. Daw, N. D. Trial-by-trial data analysis using computational models. in *Decision Making, Affect, and Learning: Attention and Performance XXIII* 1–26 (2011).

10. Sutton, R. & Barto, A. *Reinforcement Learning: An Introduction*. (MIT Press, 1998).
11. Palminteri, S., Khamassi, M., Joffily, M. & Coricelli, G. Contextual modulation of value signals in reward and punishment learning. *Nature Communications* **6**, 8096 (2015).
12. Carvalheiro, J., Conceição, V. A., Mesquita, A. & Seara-Cardoso, A. Acute stress impairs reward learning in men. *Brain and Cognition* **147**, 105657 (2021).
13. Schönberg, T., Daw, N. D., Joel, D. & O'Doherty, J. P. Reinforcement learning signals in the human striatum distinguish learners from nonlearners during reward-based decision making. *The Journal of Neuroscience* **27**, 12860–12867 (2007).
14. Zhang, L., Lengersdorff, L., Mikus, N., Gläscher, J. & Lamm, C. Using reinforcement learning models in social neuroscience: frameworks, pitfalls and suggestions of best practices. *Social Cognitive and Affective Neuroscience* **15**, 695–707 (2020).
15. Lefebvre, G., Lebreton, M., Meyniel, F., Bourgeois-Gironde, S. & Palminteri, S. Behavioural and neural characterization of optimistic reinforcement learning. *Nature Human Behaviour* **1**, 0067 (2017).
